# The LC3-interacting region of NBR1 is a protein interaction hub enabling optimal flux

**DOI:** 10.1101/2024.05.09.593318

**Authors:** Brian J North, Amelia E Ohnstad, Michael J Ragusa, Christopher J Shoemaker

## Abstract

During autophagy, potentially toxic cargo is enveloped by a newly formed autophagosome and trafficked to the lysosome for degradation. Ubiquitinated protein aggregates, a key target for autophagy, are identified by multiple autophagy receptors. NBR1 is an archetypal autophagy receptor and an excellent model for deciphering the role of the multivalent, heterotypic interactions made by cargo-bound receptors. Using NBR1 as a model, we find that three critical binding partners – ATG8-family proteins, FIP200, and TAX1BP1 – each bind to a short linear interaction motif (SLiM) within NBR1. Mutational peptide arrays indicate that these binding events are mediated by distinct overlapping determinants, rather than a single, convergent, SLiM. AlphaFold modeling underlines the need for conformational flexibility within the NBR1 SLiM, as distinct conformations mediate each binding event. To test the extent to which overlapping SLiMs exist beyond NBR1, we performed peptide binding arrays on >100 established LC3-interacting regions (LIRs), revealing that FIP200 and/or TAX1BP1 binding to LIRs is a common phenomenon and suggesting LIRs as protein interaction hotspots. Comparative analysis of phosphomimetic peptides highlights that while FIP200 and Atg8-family binding are generally augmented by phosphorylation, TAX1BP1 binding is nonresponsive, suggesting differential regulation of these binding events. In vivo studies confirm that LIR-mediated interactions with TAX1BP1 enhance NBR1 activity, increasing autophagosomal delivery by leveraging an additional LIR from TAX1BP1. In sum, these results reveal a one-to-many binding modality in NBR1, providing key insights into the cooperative mechanisms among autophagy receptors. Furthermore, these findings underscore the pervasive role of multifunctional SLiMs in autophagy, offering substantial avenues for further exploration into their regulatory functions.

## INTRODUCTION

Autophagy is a highly conserved pathway that maintains cell health by breaking down and recycling cellular components such as protein aggregates or damaged organelles. This process involves the formation of a double-membraned vesicle, the autophagosome, which encapsulates unwanted cellular cargo and transports it to lysosomes for degradation (Søreng *et al*., 2018). While autophagy occurs at a basal level to manage cell waste, cells can activate autophagy in response to urgent needs such as starvation or stress. Many stress-induced forms of autophagy are highly selective for specific cargoes. This specificity is conferred by selective autophagy receptors (SARs), proteins that identify cargo and direct it towards the autophagic machinery to ensure its efficient degradation (Kirkin and Rogov, 2019; Lamark and Johansen, 2021; Vargas *et al*., 2023).

Among these receptors, sequestrosome-1 (SQSTM1), also known as p62, is a prominent member of the SQSTM1-like receptor (SLR) family, which includes p62, NBR1, TAX1BP1, NDP52, and Optineurin. SLRs are equipped with several critical domains: ubiquitin-binding domains for cargo recognition, Atg8 interacting regions/motifs (AIMs) for attachment to autophagosomal membranes, and various domains that promote oligomerization (Johansen and Lamark, 2020). A key role of p62 is the formation of cytoplasmic clusters called p62 bodies, through a process of polymerization and phase separation (Ciuffa *et al*., 2015; Wurzer *et al*., 2015; Sun *et al*., 2018; Zaffagnini *et al*., 2018; Kurusu, Morishita and Komatsu, 2023). This mechanism is critical for the cellular clearance of protein aggregates through a specialized form of autophagy known as aggrephagy (Bauer, Martens and Ferrari, 2023).

The SLR family exhibits significant functional interconnectivity, with both unique and overlapping roles that enable a coordinated response to diverse cellular challenges. Although p62 is often highlighted in studies of SLR function, NBR1 is considered an ancestral SLR with a gene duplication in early metazoans giving rise to p62 (Svenning *et al*., 2011; Rasmussen *et al*., 2022). This evolutionary split allowed NBR1 and p62 to evolve distinct functions despite their structural similarities. For example, p62 retains an extended PB1 domain essential for forming p62 filaments, while NBR1, having lost one binding face of its PB1 domain, depends on interaction with the p62 PB1 domain for filament recruitment. (Lamark *et al*., 2003; Wilson *et al*., 2003; Ciuffa *et al*., 2015; Jakobi *et al*., 2020). NBR1 recruitment is thought to cap the p62 filaments and enhance binding to polyubiquitinated substrates through its higher-affinity UBA domain (Turco *et al*., 2021). Following the sequestration of ubiquitinated cargo by p62 and NBR1, TAX1BP1 is recruited to p62 bodies through its interaction with NBR1. Together, TAX1BP1 and NBR1 recruit FIP200, a crucial autophagy initiation factor, thereby initiating autophagosome formation around cargo (Ohnstad *et al*., 2020; Turco *et al*., 2021). This collaborative action among SLRs promotes efficient encapsulation and degradation of cellular debris, underscoring their collective role in maintaining cellular health.

The functionality of SARs is also closely linked to their interaction with the autophagosomal membrane facilitated by SAR binding to mammalian Atg8-family proteins, which include the LC3 and GABARAP subfamilies. This interaction is mediated by short linear interaction motifs (SLiMs) known as Atg8-family interacting motifs (AIMs) or LC3-interacting regions (LIRs) (Birgisdottir, Lamark and Johansen, 2013). LIRs have emerged as a key focus of autophagy research. These motifs, with their consensus sequence F/W/Y-X-X-L/I/V, are typically embedded within intrinsically disordered regions and lack stable 3D structures in isolation (Popelka and Klionsky, 2015). However, they can adopt specific, constrained conformations when binding to their targets (Rogov *et al*., 2023). In addition, some SARs feature a FIP200-interacting region (FIR) that can resemble, and sometimes co-exist with, the LIR (Turco *et al*., 2019; Fu *et al*., 2021; Zhou *et al*., 2021; Goodall, Kraus and Harper, 2022). While these interactions are typically of modest affinity, LIR and FIR binding can be significantly enhanced by post-translational modifications such as phosphorylation, further underscoring the complex regulation of autophagy receptors (Gubas and Dikic, 2022).

In this study, we investigate the role of multivalent, heterotypic interactions in selective autophagy using the archetypal receptor NBR1. We demonstrate that NBR1 binds three key partners–Atg8-family proteins, FIP200, and TAX1BP1–through a single region containing a short SLiM. Expanding our analysis to over 100 LIR-containing peptides, we observe frequent overlap between LC3-, FIP200-, and TAX1BP1-interacting regions (TIRs). Mutational peptide arrays indicate that each binding event is governed by unique molecular determinants, rather than a shared motif. Complementary in vivo studies reveal that while SLRs typically compete, TAX1BP1 uniquely enhances NBR1 flux, reliant on NBR1’s multifunctional SLiM. These findings underscore the organizational complexity of this one-to-many binding modality, providing detailed insights into the cooperative dynamics among autophagy receptors and emphasizing the critical, versatile role of multifunctional SLiMs in regulating autophagy.

## RESULTS

### NBR1 contains overlapping SLiMs that recruit TAX1BP1, LC3A, and FIP200

NBR1 is an archetypal autophagy receptor and an excellent model for deciphering the role of multivalent, heterotypic interactions made by cargo-bound receptors (Rasmussen *et al*., 2022). NBR1, TAX1BP1, and p62 were previously reported to interact, which enhances the process of aggrephagy (Kirkin, Lamark, Johansen, *et al*., 2009; Kirkin, Lamark, Sou, *et al*., 2009; Zaffagnini *et al*., 2018; Sánchez-Martín *et al*., 2020; Schlütermann *et al*., 2021; Turco *et al*., 2021; Vargas *et al*., 2023). In vitro, the interaction between NBR1 and TAX1BP1 was reported to depend on a conserved FW domain of NBR1 (Turco *et al*., 2021). To corroborate this interaction and investigate whether other NBR1 domains also contribute to binding TAX1BP1 in vivo, we systematically truncated NBR1 from both its N- and C-termini (**Fig S1**). We co-transfected HEK293T cells with myc-tagged TAX1BP1 and TagBFP-V5-tagged NBR1 variants. We then immunoprecipitated NBR1 and monitored for co-immunoprecipitation of myc-TAX1BP1 (**Fig 1A**). Among these truncations, NBR1^1-622^ and NBR1^781-end^ emerged as the longest N- and C-terminal fragments, respectively, lacking observable TAX1BP1 binding, suggesting the presence of a TAX1BP1 binding site in this range (i.e. NBR1^623-780^) (**Fig 1A**). Intriguingly, this region does not contain the FW domain but contains two other notable motifs: coiled-coil 2 (CC2, residues 686-727), previously reported to bind FIP200 (Turco *et al*., 2021), and a well-characterized short linear interaction motif (SLiM), the LC3-interacting region (LIR, residues 732-735) (Kirkin, Lamark, Sou, *et al*., 2009).

**Figure 1.**
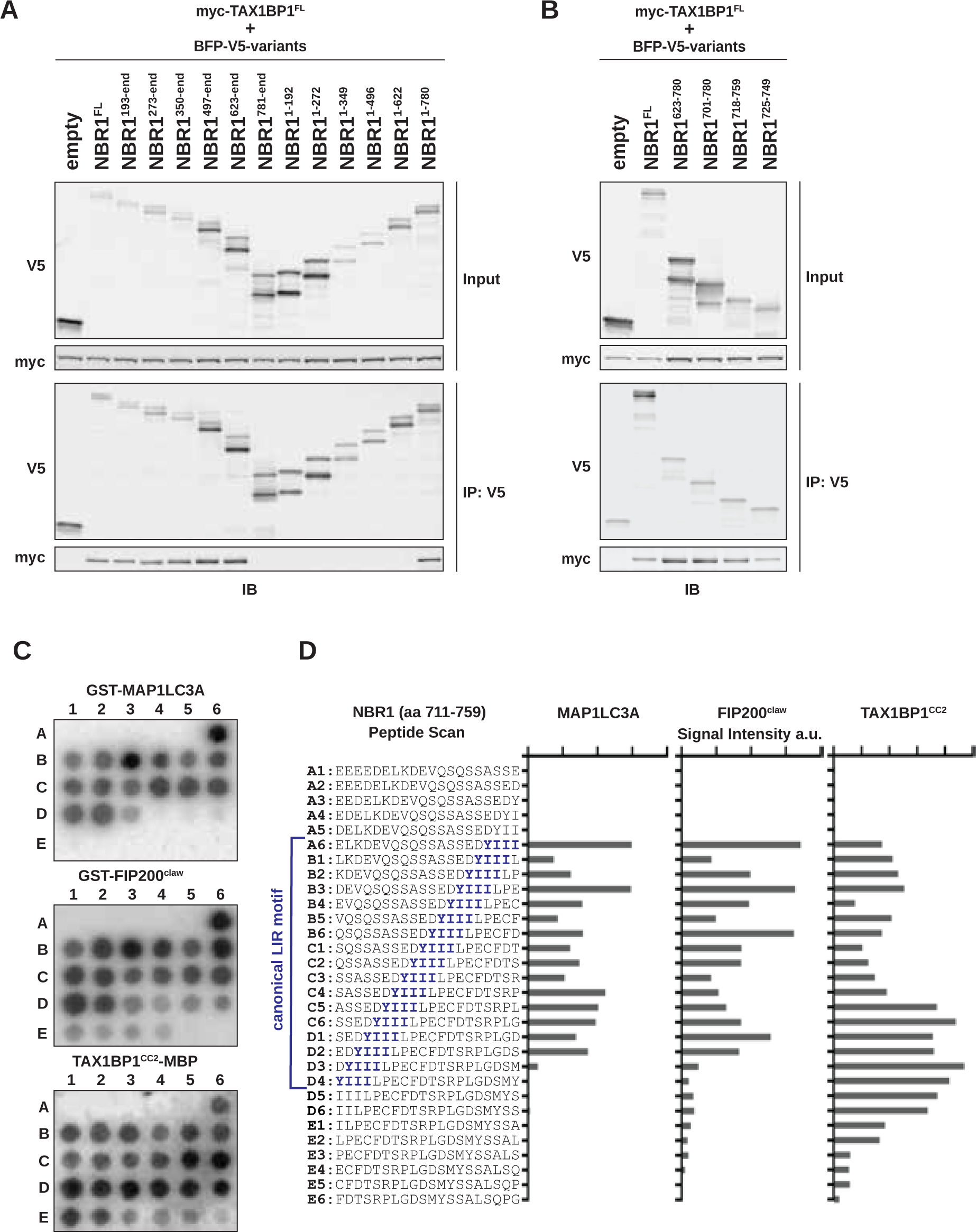
A short unstructured region in NBR1 mediates interactions with MAP1LC3A, FIP200, and TAX1BP1. **A, B)** HEK293T cells were co-transfected with full-length (FL) myc-TAX1BP1 and indicated BFP-V5-NBR1 truncations. Extracts derived from transfected cells were immunoprecipitated (IP) with protein G dynabeads conjugated with anti-V5 antibody. Input and eluates were resolved by SDS-PAGE followed by immunoblotting (IB) with the indicated antibodies. **C)** A sliding frame peptide array spanning NBR1^711-759^ was probed with 100 nM GST-MAP1LC3A, GST-FIP200^Claw^ (residues 1490-1594), or TAX1BP1^CC2^-MBP (residues 346-506). Antibody-HRP conjugates were used to visualize binding to the arrays. Each spot represents a 1 residue register shift. **D)** Quantification of HRP-derived intensities from C. Peptide sequences and corresponding positions in C are indicated (A1-E6). The canonical NBR1 LIR motif is indicated in blue. a.u., arbitrary units.

To pinpoint the NBR1-TAX1BP1 interaction site, we made additional truncations of the NBR1^623-780^ region, identifying NBR1^725-749^ as sufficient for interaction with TAX1BP1 (**Fig 1B**). This segment harbors the NBR1 LIR (Rozenknop *et al*., 2011), suggesting a potential overlap between the TAX1BP1-interacting region (TIR) and LC3-interacting region. Notably, two peptides that failed to pulldown TAX1BP1 give important clues to a potential TIR motif (**Fig S1B**). NBR1^735-754^ lacks the LIR and NBR1^730-744^ contains the LIR but lacks downstream residues, yet neither peptide was sufficient for binding, suggesting a related but distinct mode of binding between NBR1 and TAX1BP1 or LC3-family proteins. As a control, we confirmed NBR1 LIR functionality using GST-tagged LC3A as bait, confirming NBR1^725-749^ is sufficient to interact with LC3A (**Fig. S1C**). Thus, the extended LIR of NBR1 plays a dual role in binding TAX1BP1 and Atg8-family proteins.

Intriguingly, within the initially identified TAX1BP1-binding region of NBR1^623-780^ lies CC2, previously recognized for its interaction with the Claw domain of FIP200 (Turco *et al*., 2021). To validate this interaction, we purified a GST-tagged FIP200 Claw domain (1490-1594, hereafter FIP200^Claw^), immobilized it on GST-resin, and probed with HEK293T cell lysates expressing various NBR1 truncations. As anticipated, removal of CC2 led to reduced NBR1 association with FIP200^Claw^, consistent with previous findings (Turco *et al*., 2021), yet NBR1^725-749^ maintained a reduced FIP200^Claw^ interaction (**Fig S1D**). As such, a single unstructured region of NBR1, residues 725-749, mediates the interaction between NBR1 and an array of critical autophagy mediators.

To further characterize these binding events, we used fluorescence anisotropy to determine the relative affinity of these interactions using a fluorescently-labelled NBR1 peptide (FITC-NBR1^725-749^). LC3A bound with a K_D_ of 1.79 ± 0.17 μM, consistent with previous literature (Rozenknop *et al*., 2011). FIP200^Claw^ bound with a K_D_ of 16.66 ± 3.22 μM. TAX1BP1 coiled-coil 2 (residues 346-506, hereafter TAX1BP1^CC2^), which we previously reported to be required for TAX1BP1 binding to NBR1, bound with a K_D_ of 5.82 ± 0.66 μM (**Fig S1E**) (Ohnstad *et al*., 2020). Taken together, all three binding events occur with micromolar affinity and well within the range of previously reported affinities for LC3- and FIP200-interacting proteins (Wirth *et al*., 2019, 2021; Zhou *et al*., 2021).

Three binding interactions mediated by a short, unstructured span of NBR1 implies that its interaction with FIP200 and TAX1BP1, akin to LC3A, are orchestrated by SLiMs. To precisely map these motifs, we constructed a single amino acid sliding-frame peptide array spanning NBR1 residues 711-759. We probed the sliding frame peptide array with purified GST-tagged LC3A, GST-tagged FIP200^Claw^, or MBP-tagged TAX1BP1^CC2^ and visualized with HRP-conjugated antibodies targeting GST or MBP (**Fig 1C**, **Fig 1D**). LC3A binding correlated strongly with the presence of the core LIR motif (YIII, spots A6-D3), validating our approach. Similarly, FIP200^Claw^ primarily engaged the core LIR. In contrast, maximal binding of TAX1BP1^CC2^ did not fully align with the LIR itself but with subsequent residues (spots C5-D6), suggesting an extended or alternative motif beyond the core LIR motif (**Fig 1D**).

### FIR and TIR overlap with LIRs is common despite divergent motifs

Multiple instances of LIRs overlapping with FIRs have been demonstrated (Turco *et al*., 2019; Zhou *et al*., 2021; Goodall, Kraus and Harper, 2022), while other examples of FIRs that are not LIRs have also been documented (Fujita *et al*., 2013; Smith *et al*., 2018; Ravenhill *et al*., 2019; Fu *et al*., 2021; Zhou *et al*., 2021). This raises a fundamental question: To what extent do LIRs and FIRs overlap? Moreover, our demonstration of TAX1BP1 binding in relation to a LIR and FIR prompts an additional question: do LIRs or FIRs also bind TAX1BP1? To answer these questions, we printed a peptide array comprising 100 established LIR sequences from the LIR-Central database (Chatzichristofi *et al*., 2023) (**Supplemental Table 1**). As a control, we included the multifunctional NBR1^725-749^ sequence identified in Fig 1B. We also included 5 previously reported FIR peptides as a control for FIP200 binding (Fujita *et al*., 2013; Smith *et al*., 2018; Ravenhill *et al*., 2019; Fu *et al*., 2021; Zhou *et al*., 2021).

The LIR family array was probed as in Fig 1C by incubating with purified HA-tagged LC3A, HA-tagged GABARAPL1, GST-tagged FIP200^Claw^, or MBP-tagged TAX1BP1^CC2^. Each protein probe was subsequently visualized via HRP-conjugated antibody. Binding of LC3A, GABARAPL1, FIP200^Claw^ and TAX1BP1 was prevalent, but not universal. LC3A bound 29 peptides, GABARAPL1 bound 49 peptides, FIP200 bound 57 peptides, and TAX1BP1 bound 49 peptides (**Fig 2A and B**). These data show that among the spectrum of known LIR-containing peptides, there exists an equally rich spectrum of FIR motifs and an unexplored array of potential TIR motifs. In comparison, previously validated FIR peptides were largely selective for FIP200^Claw^ binding (**Fig 2B**). We note that 32 LIR peptides never bound to any of the four probes. As we cannot distinguish true non-binding events (e.g. spurious LIRs) from technical artifacts (e.g. peptide aggregation or unsuccessful peptide printing), “never-binders” were excluded from further analysis (**Fig 2A**, red asterisks).

**Figure 2.**
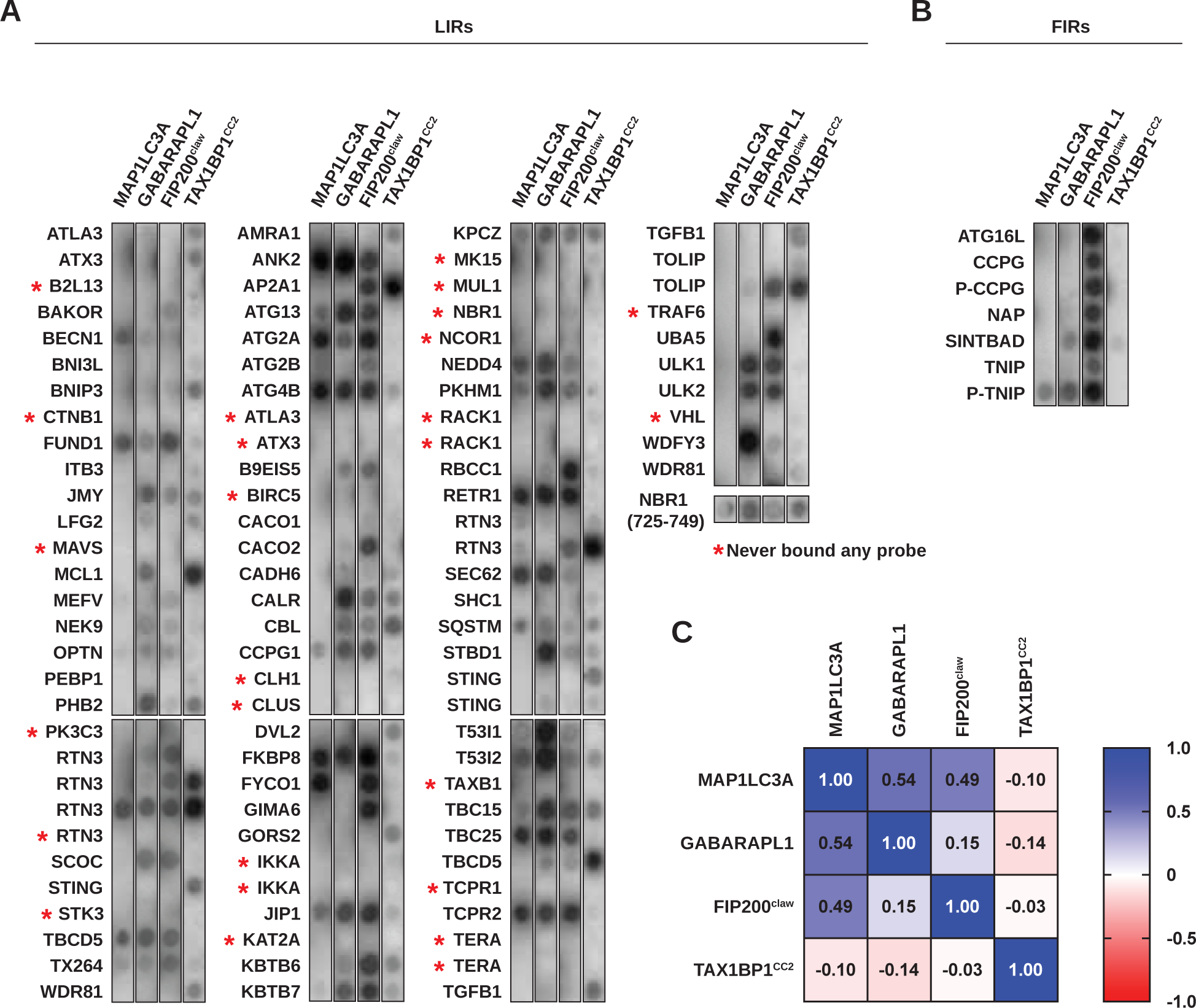
Comprehensive analysis of LIR binding interactions with Atg8-family proteins, FIP200^Claw^, and TAX1BP1^CC2^. **A)** The LIR peptide array, containing peptides selected from the iLIR database, was probed with 100 nM GST-LC3A, GST-GABARAPL1, GST-FIP200^Claw^, or TAX1BP1^CC2^-MBP. Antibody-HRP conjugates were used for visualization. Peptide spots that bound none of four protein probes (“never-binders”) are labelled with red asterisk and excluded from further analysis. Redundant protein names reflect multiple LIR sequences within a single protein. **B)** FIR containing peptides from selected literature, probed as in A. **C)** Pearson correlation analysis of SLiM binding preference for each protein (LC3A, GABARAPL1, GST-FIP200^Claw^, or TAX1BP1^CC2^-MBP) based on spot intensity in A and B. “Never-binders” and phosphomimetic peptides were exclude from analysis. *N* = 79.

To assess the relationship between LC3-interacting regions, FIP200-interacting regions, and TAX1BP1-interacting regions, we conducted a correlation analysis based on binding intensity (**Fig 2C**). Our analysis reveals that the binding of LC3A and GABARAPL1 is moderately correlated (*r* = 0.54, *p*=<0.001, *N*=79), reflecting the previously reported preference of individual LIR motifs for different Atg8-family proteins (Rogov *et al*., 2017; Atkinson *et al*., 2019; Wirth *et al*., 2019). Similarly, LC3A-binding and FIP200^Claw^-binding are moderately correlated (*r* = 0.49, *p*=<0.001), suggesting related motif elements between FIRs and LIRs. In contrast, the correlation between GABARAPL1-binding and FIP200-binding is weak (*r* =.147, *p*=.19), highlighting that not all LIR motifs are FIR motifs and that LC3A binding is a better predictor of FIR motifs than GABARAPL1 binding. Furthermore, TAX1BP1-binding showed no significant correlation with LC3A binding (*r*=-0.09, *p*=.39), GABARAPL1 binding (*r*=-0.13*, p*=.22), or FIP200 binding (*r*=-0.03, *p*=.79). These results suggest that the mechanism of TAX1BP1-binding is distinct from that of the core LIR motif.

### Related, but distinct, binding motifs mediate LC3A, FIP200, and TAX1BP1 binding

Discrepancies in binding among LC3A, FIP200, and TAX1BP1 led us to explore the specific binding determinants recognized by these proteins. To this end, we created a mutational peptide array for the minimally sufficient NBR1^725-749^ peptide, systematically mutating each amino acid position to all other amino acids. The array was probed as in Fig 1C using HA-LC3A, GST-FIP200^Claw^, or MBP-TAX1BP1^CC2^. As expected, LC3A binding required a consensus F/W/Y-X-X-L/I/V motif, with additional contributions from negatively charged sites at positions −2, −1, and +6 relative to the first residue of the LIR (**Fig 3A** and **3D**). Interestingly, the FIP200^Claw^ probe revealed high tolerance to single point mutations within the core hydrophobic motif of NBR1, yet was sensitive to the removal of acidic residues from the peptide (i.e. positions −2, −1, +6, +7) (**Fig 3B** and **3D**). In contrast, TAX1BP1^CC2^ displayed a distinct binding pattern, relying on residues at the +4, +5, and +8 positions relative to the core LIR motif (**Fig 3C** and **3D**). These findings collectively support that LC3A, FIP200^Claw^, and TAX1BP1^CC2^ interact through unique yet overlapping short linear motifs within an unstructured region of NBR1.

**Figure 3.**
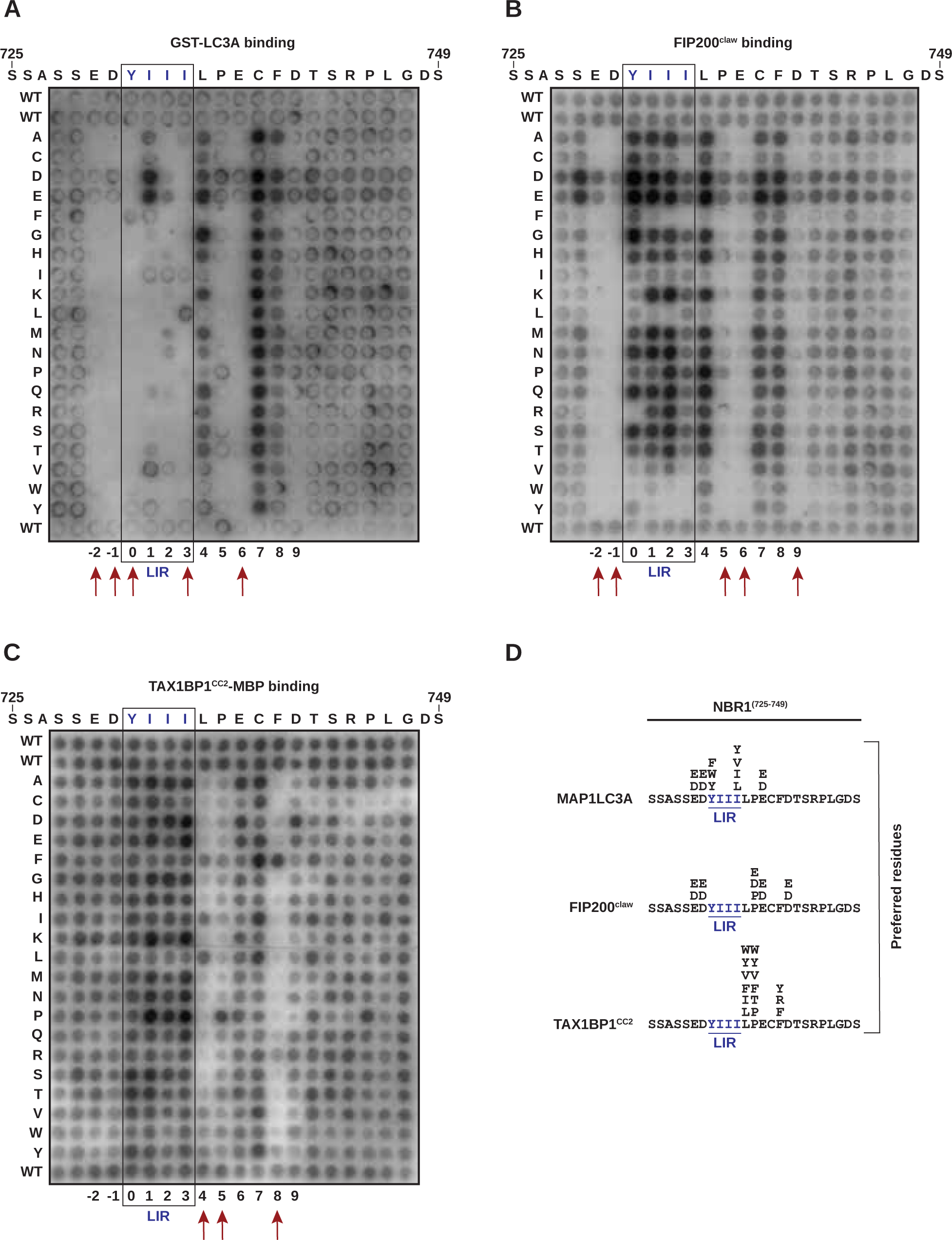
Mutational analysis of NBR1 reveals unique binding determinants for MAP1LC3A, FIP200^Claw^, and TAX1BP1^CC2^. **A-C)** Mutational peptide array of NBR1^728-747^ probed with GST-LC3A (**A**), GST-FIP200^Claw^ (**B**), or TAX1BP1^CC2^-MBP (**C**) and immunobloted with HRP-conjugated antibodies to GST or MBP. Each amino acid position was substituted for every other amino acid. NBR1’s canonical LIR motif (YIII) is indicated in blue. Red arrows indicate key residues identified for each SLiM binding motif. **D)** Summary of preferred residues inferred from **A-C**. Core LIR motif labelled in blue.

Phosphorylation has been previously reported to enhance binding to multiple LIRs and FIRs (Rogov *et al*., 2023). Given the notable contribution of acidic residues to NBR1 binding by LC3A and FIP200, we aimed to more systematically evaluate the contribution of phosphorylation to Atg8-family and FIP200 binding. Noting the importance of phosphorylation at the −2 or −1 positions, we identified all LIRs from our array containing possible phosphorylation sites (serine or threonine residues) at these positions (**Supplemental Table S1**). We then mutated each residue to aspartic acid and evaluated the effect of the phosphomimetic residues on the interaction with Atg8-family proteins (both LC3A and GABARAPL1), FIP200, and TAX1BP1 (compare **Fig 2A**, column 1, and **Fig 4A**). LC3A and GABARAPL1 showed a significant increase in signal intensity with phosphomimetic variants (*p*=0.054 and *p*=0.0042, respectively), as did FIP200^Claw^ (*p*=< 0.0001), aligning with previous findings (**Fig 4B**) (Rogov *et al*., 2023). However, among TIR-containing peptides, TAX1BP1^CC2^ did not display a clear preference for phosphomimetic variants (**Fig 4B**).

**Figure 4.**
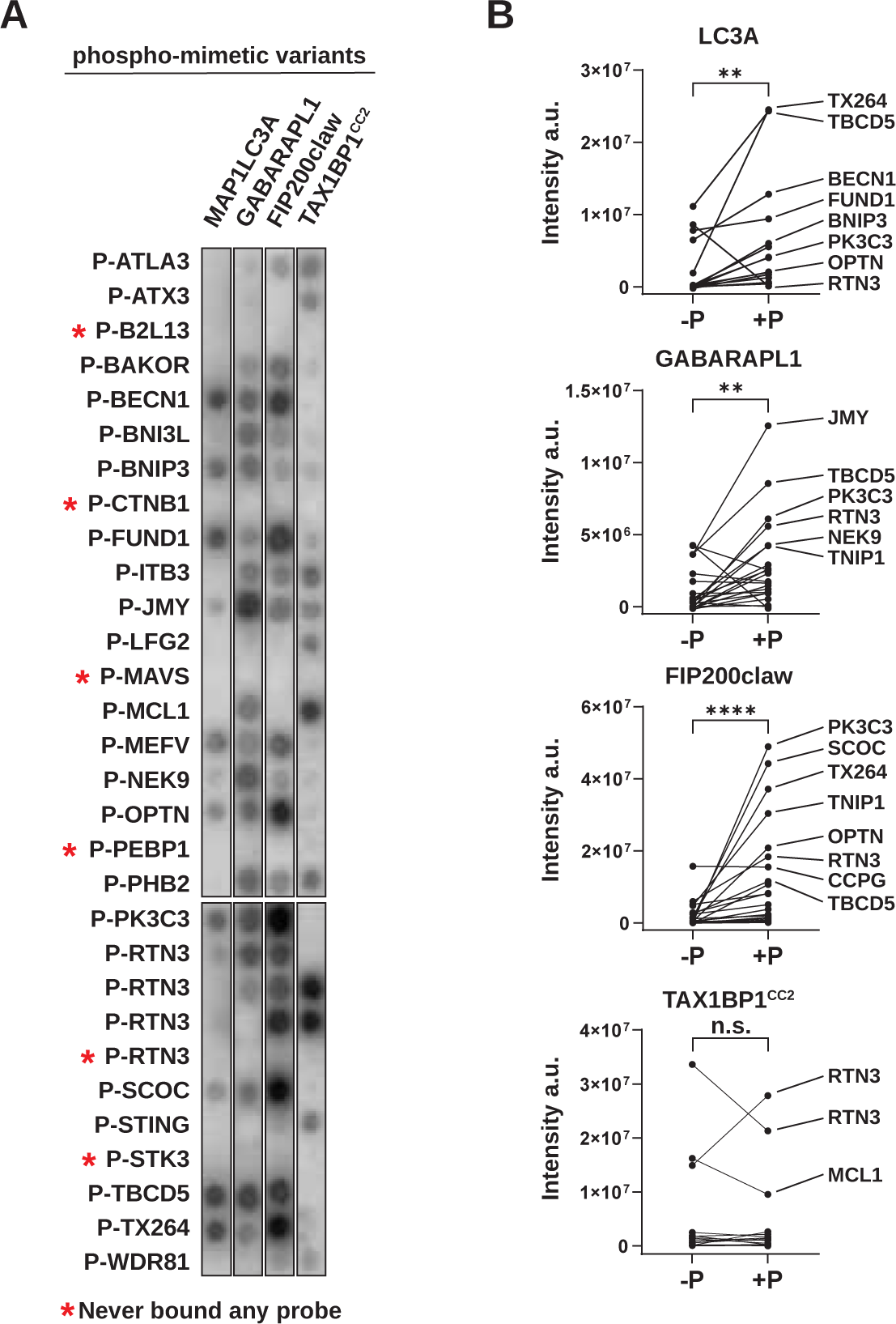
Phosphorylation differentially regulates LIRs, FIRs, and TIRs. **A)** Phosphomimetic peptide variants were generated for each LIR peptide containing phosphorylatable residues (Ser or Thr) at the −2 or −1 positions relative to the core LIR motif. The phospho-LIR peptide array was probed with 100 nM GST-LC3A, GST-GABARAPL1, GST-FIP200^Claw^, or TAX1BP1^CC2^-MBP. The phosphomimetic LIR array was probed and imaged simultaneously with the native LIR array in Fig. 2A to allow comparison. Antibody-HRP conjugates were used to visualize the arrays. Peptide spots that bound none of four protein probes (“never-binders”) are labelled with red asterisk and excluded from further analysis. **B)** Comparison of spot intensity between native (Fig. 2A) and phosphomimetic variant peptides (A) using a nonparametric t-test with Wilcoxon matched pairs signed rank test. **, *p*<0.01; ****, *p*<0.0001; *ns*, not significant.

To further characterize TAX1BP1 binding, we focused on the contribution of the +4, +5, and +8 positions within NBR1^725-749^. Co-immunoprecipitation of full-length myc-TAX1BP1 and BFP-V5-tagged NBR1 variants (L736A, P737A, and F740A) confirmed that these positions contribute to TAX1BP1 binding relative to a Y732A LIR mutant (**Fig 5A and Fig S2A**) (Kirkin, Lamark, Sou, *et al*., 2009). Conversely, the F740A mutation did not impair FIP200^Claw^ or LC3A binding as compared to the NBR1^Y732A^ control, indicating its specificity for TAX1BP1 despite its proximity to the LIR (**Fig S2B** and **S2C**). Thus, Y732A and F740A mutations can be used to selectively disrupt interactions with LC3A and TAX1BP1, respectively.

**Figure 5.**
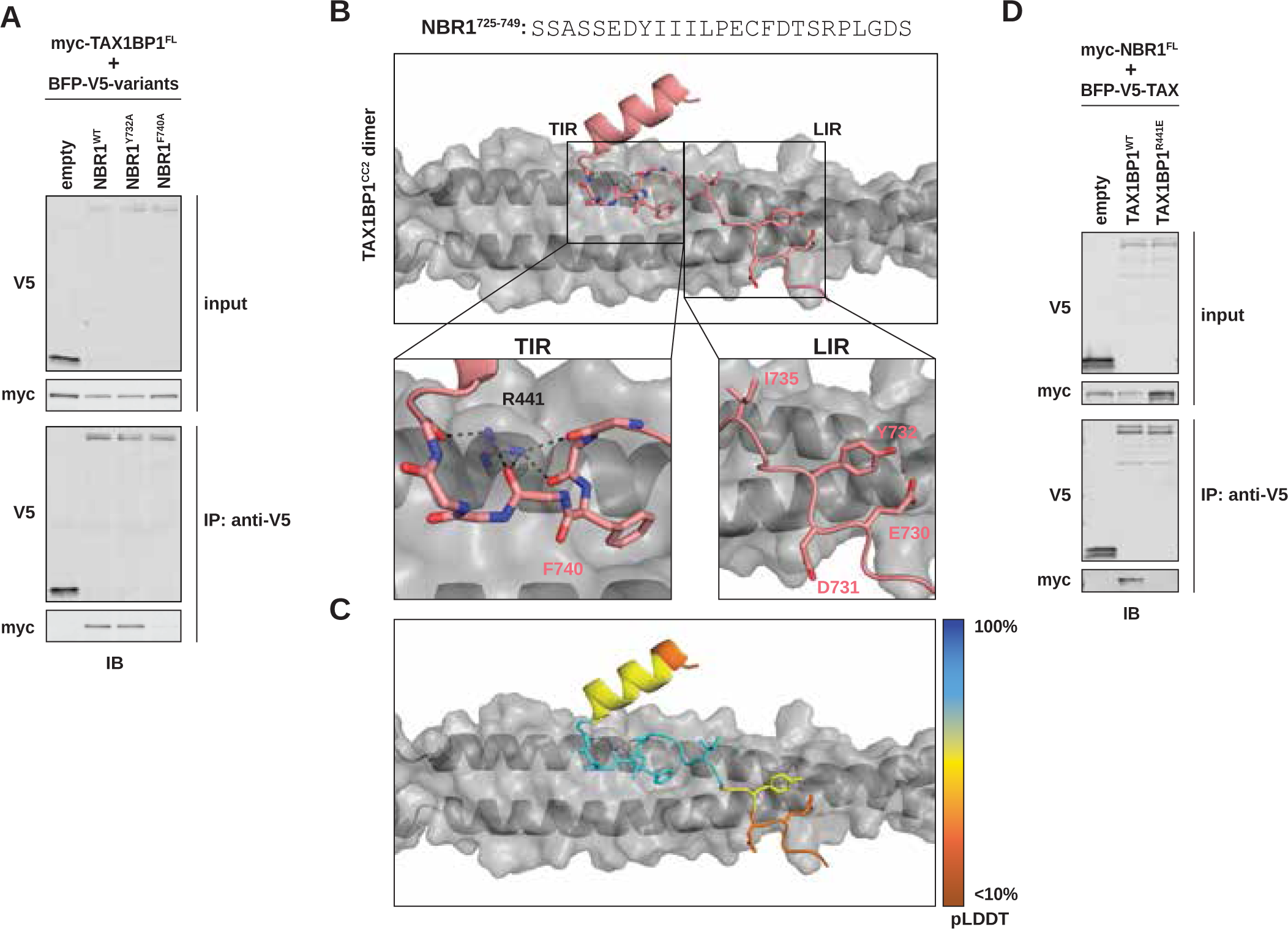
Structural modeling reveals a unique interface between TAX1BP1 and NBR1. **A)** HEK293T cells were co-transfected with full-length (FL) myc-TAX1BP1 and indicated BFP-V5-NBR1^FL^ variants. Extracts derived from transfected cells were immunoprecipitated (IP) with protein G dynabeads conjugated with anti-V5 antibody. Input and eluates were resolved by SDS-PAGE followed by immunoblotting (IB) with the indicated antibodies. **B)** Predictive structural model of NBR1^725-749^ with a dimer TAX1BP1^CC2^. Model was generated using AlphaFold AF2-multimer with MMseqs2 through Google Colab. The TIR inset highlights the hydrophobic binding pocket occupied by NBR1^F740^ and the backbone interactions mediated by TAX1BP1^R441^. The LIR inset highlights the lack of meaningful interactions between NBR1^LIR^ and TAX1BP1^CC2^. **C)** Predicted IDDT per position (pLDDT) for NBR1^725-749^ as an estimation of model confidence. Blue is high confidence, orange is low confidence. **D)** HEK293T cells were co-transfected with full-length (FL) myc-NBR1 and indicated BFP-V5-TAX1BP1 variants. Extracts derived from transfected cells were immunoprecipitated (IP) with protein G dynabeads conjugated with anti-V5 antibody. Input and eluates were resolved by SDS-PAGE followed by immunoblotting (IB) with the indicated antibodies.

### Structural modeling identifies a unique binding mode between NBR1 and TAX1BP1

Numerous structural studies have detailed the interaction between LIRs and Atg8-family proteins, as well as the interaction between FIRs and FIP200^Claw^ (Fu *et al*., 2021; Zhou *et al*., 2021; Rogov *et al*., 2023). However, the structural basis of the NBR1-TAX1BP1 interaction was uncharacterized. To better understand how NBR1 interacts with TAX1BP1, we utilized AlphaFold structural modeling to predict the interface between NBR1 and TAX1BP1 and compared it to the predicted NBR1 interactions with LC3A and FIP200^Claw^ (Mirdita *et al*., 2022).

Our AlphaFold model, pairing NBR1^725-749^ with a dimer of TAX1BP1^CC2^, provides several key insights. Notably, while the NBR1 LIR motif makes high-confidence interactions with LC3A and FIP200^Claw^ (**Fig S3A-F**), it makes minimal contribution to TAX1BP1 binding (**Fig 5B** (LIR inset) and **5C**). Instead, residue F740 of NBR1 embeds in a hydrophobic pocket on either side of the TAX1BP1 coiled-coil dimer (**Fig 5B** (TIR inset), a finding supported by our peptide array and co-immunoprecipitation data, which independently identified F740 as a crucial determinant of NBR1 binding to TAX1BP1.

To further assess our structural predictions we performed buried surface area (BSA) analysis using the PDBePISA server (**Fig S3G**) (Krissinel and Henrick, 2007). This analysis highlights the largely overlapping motifs recognized by FIP200 and LC3A. In addition, the BSA analysis emphasizes the importance of distinct amino acids in NBR1 for its interaction with TAX1BP1, especially F740 which has a BSA of 104.62 Å^2^ upon binding to TAX1BP1 but a BSA of 0 Å^2^ upon binding to either LC3A or FIP200.

Moreover, our structural model predicts that Arg441 of TAX1BP1 forms a finger-like protrusion that makes backbone interactions with the +6, +7, +9, and +12 positions of the NBR1 peptide, inducing a small kink in NBR1 to accommodate binding (**Fig 5B** (TIR inset)). To validate this model, we generated a charge inversion mutant, TAX1BP1^R441E^. Co-transfection of HEK293T cells with full-length myc-tagged NBR1 and BFP-V5 tagged TAX1BP1 variants showed that R441E inhibits the interaction of TAX1BP1 with NBR1 (**Fig 5D**). This finding underscores the predictive power of our model and demonstrates that disruption of either side of the binding interface, NBR1^F740A^ or TAX1BP1^R441E^, is sufficient to inhibit the interaction between full length NBR1 and TAX1BP1. Moreover, these effects are specific as the TAX1BP1^R441E^ mutation does not affect TAX1BP1 dimerization, another critical facet of TAX1BP1^CC2^ (**Fig S3H**). Together, these data show that varied conformations taken by a short, disordered region of NBR1 bring key residues into contact with diverse binding pockets on LC3A, FIP200^Claw^, and TAX1BP1^CC2^.

### NBR1 recruits TAX1BP1 to enhance its own lysosomal delivery

What effect does the one-to-many interaction of the NBR1 SLiM have in vivo? Dissecting individual points of connection among SQSTM1-like receptors (SLRs) is complex due to their interconnectivity. To overcome this, we utilized a previously characterized HeLa cell line lacking all five major SLRs: p62, NBR1, TAX1BP1, NDP52, and Optineurin (PentaKO cells) (Lazarou *et al*., 2015). To monitor lysosomal delivery in these cells, we employed a suite of tandem fluorescent (tf) reporters composed of RFP and GFP fused to the N-terminus of each cargo receptor (Shoemaker *et al*., 2019). In the acidic environment of the lysosome, GFP fluorescence is quenched, while the RFP signal remains, making the red:green fluorescence ratio a reliable indicator of lysosomal delivery (**Fig 6A**).

**Figure 6.**
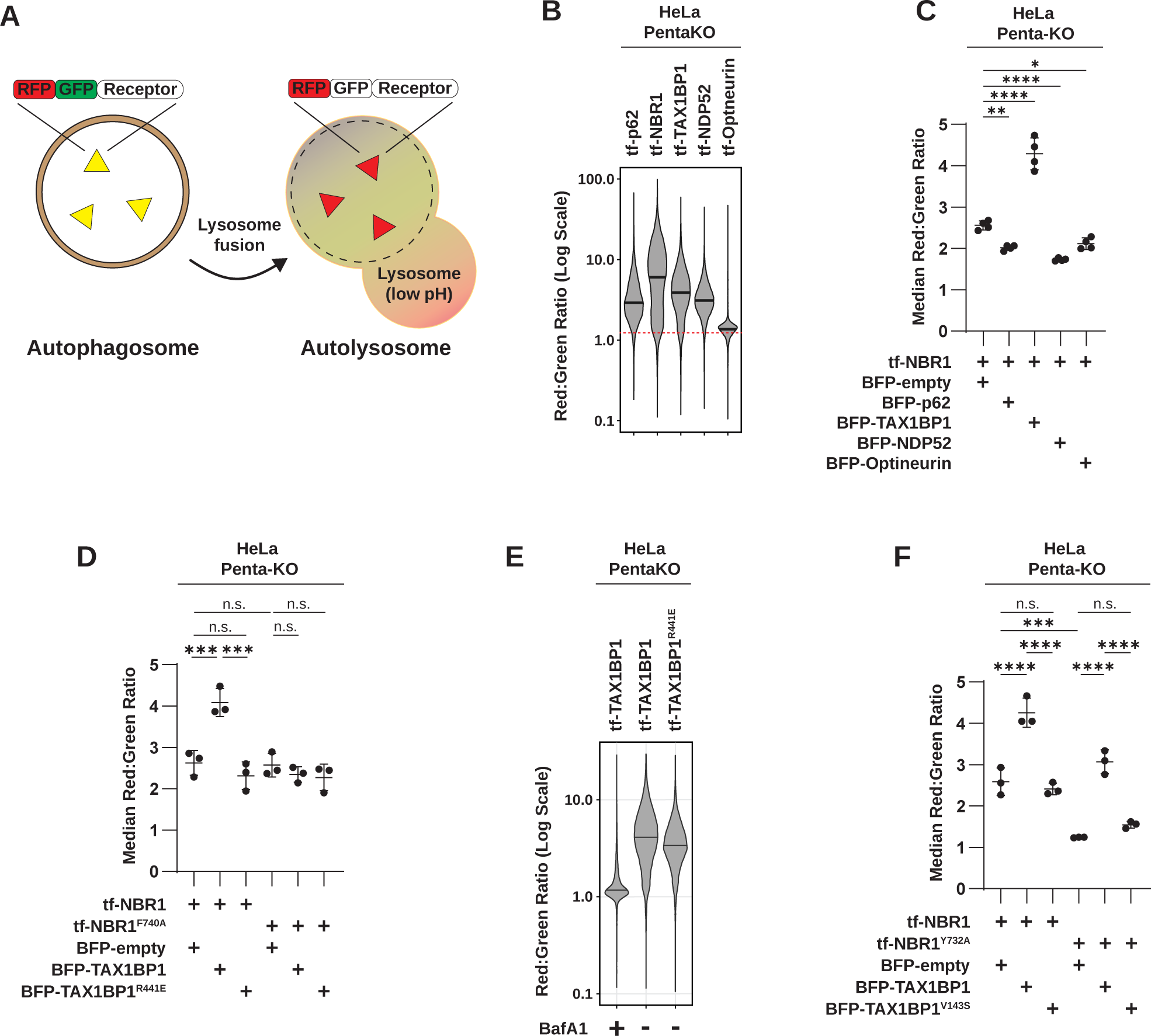
Synergy between NBR1 and TAX1BP1 mitigates competition. **A)** Schematic of the tandem fluorescent (tf) reporter system. Throughout autophagosome formation, both RFP and GFP fluorescence are observed. Upon lysosomal fusion, the acidic lysosomal environment quenches GFP, whereas RFP is maintained. Therefore, increasing red:green fluorescence ratio corresponds with increased lysosomal delivery. **B)** HeLa PentaKO cells were transfected with tf-p62, tf-NBR1, tf-TAX1BP1, tf-NDP52, or tf-Optineurin. After 18h, cells were analyzed for red:green ratio by flow cytometry. Median values for each sample are identified by a black line within each violin. The red doted line across all samples corresponds to red:green ratio of maximally inhibited conditions (e.g. BafA1 treatment) *n* > 10,000 cells per sample. **C)** HeLa PentaKO cells were co-transfected with tf-NBR1 and indicated BFP-tagged receptors. After 18h, cells were analyzed for red:green ratio by flow cytometry. Graphs represent mean +/- SD from four independent experiments. *n* > 10,000 cells per sample. *p* values were determined using a one-way ANOVA (*p* = <0.0001). Multiple comparisons were made using Dunnet T3 correction. *, *p* < 0.05; **, *p* < 0.01; ***, *p* < 0.001; *ns*, not significant. **D)** HeLa PentaKO cells were co-transfected with the indicated tf-NBR1 and BFP-TAX1BP1 variants. After 18h, cells were analyzed for red:green ratio by flow cytometry (*n* > 10,000 cells per sample). Graphs represent mean +/- SD from three independent experiments. *p* values were determined using a one-way ANOVA (*p* = < 0.0001). Multiple comparisons were made using Tukey correction. ***, *p* < 0.001; *ns*, not significant. **E)** HeLa pentaKO cells were transfected with tf-TAX1BP1 (WT or R441E). After 18h, cells were analyzed for red:green ratio by flow cytometry (*n* > 10,000 cells per sample). Median values for each sample are identified by a black line within each violin. BafA1, Bafilomycin A1. **F)** HeLa pentaKO cells were co-transfected with tf-NBR1 (WT or Y732A LIR mutant) and BFP-TAX1BP1 (WT or V143S LIR mutant). After 18h, cells were analyzed for red:green ratio by flow cytometry (*n* > 10,000 cells per sample). Graphs represent mean +/- SD from three independent experiments. *p* values were determined using a one-way ANOVA (*p* = < 0.0001). Multiple comparisons were made using Tukey correction. ***, *p* < 0.001; ****, *p* < 0.0001; *ns*, not significant.

To validate this approach, we transfected each tf-labelled receptor into PentaKO cells and evaluated its lysosomal delivery by flow cytometry. We observed substantial flux in all reporters except tf-Optineurin, with tf-NBR1 exhibiting the highest flux (**Fig 6B**). Co-transfection with BFP-tagged p62, NDP52, or Optineurin modestly reduced tf-NBR1 flux (*p*=0.0043, *p*=<0.0001, *p*=0.018, respectively) relative to BFP alone, indicative of competitive interactions among the receptors. In contrast, co-transfection with BFP-TAX1BP1 substantially increased tf-NBR1 flux (*p*=<0.0001), demonstrating that TAX1BP1 acts synergistically with NBR1 to boost its lysosomal delivery (**Fig 6C**). This finding underscores the unique role of TAX1BP1 in enhancing NBR1-mediated autophagic pathways in vivo.

Given the previously reported interactions between p62, NBR1, and TAX1BP1 in p62 bodies, we investigated whether addition of p62 to the NBR1:TAX1BP1 complex would further enhance NBR1 flux. We co-transfected PentaKO HeLa cells with tf-NBR1, HALO-TAX1BP1 and/or BFP-p62 and examined the red:green ratio via flow cytometry. Consistent with earlier findings (**Fig 6C**), addition of BFP-p62 modestly decreased tf-NBR1 flux, while HALO-TAX1BP1 significantly increased it (**Fig S4A**). The combined addition of BFP-p62 and HALO-TAX1BP1 was largely additive, with p62 muting the TAX1BP1-mediated enhancement of tf-NBR1 flux, suggesting that p62 is unexpectedly inhibitory in our add-back system (see Discussion for details).

To rigorously test whether the synthetic enhancement of tf-NBR1 by TAX1BP1 depends on NBR1:TAX1BP1 binding, we generated tf-NBR1^F740A^ and BFP-tagged TAX1BP1^R441E^ variants, which each disrupt the NBR1:TAX1BP1 interaction. Strikingly, we found no difference in flux between tf-NBR1 and tf-NBR1^F740A^ or tf-TAX1BP1 and tf-TAX1BP1^R441E^, indicating that neither mutation affects receptor flux individually (**Fig 6D** and **6E**). However, co-transfection of PentaKO HeLa cells with tf-NBR1 (or tf-NBR1^F740A^) and BFP-empty, BFP-TAX1BP1, or BFP-TAX1BP1^R441E^ revealed that either binding mutant (NBR1^F740A^ or TAX1BP1^R441E^) was sufficient to block the enhancement of NBR1 flux (**Fig 6D**). This observation was further corroborated in Atg7-deficient K562 cells, where NBR1 degradation depends on its interaction with TAX1BP1 (Ohnstad *et al*., 2020). Transfection of ATG7^KO^/TAX1BP1^KO^ cells with wild-type TAX1BP1 rescued NBR1 degradation, whereas the R441E mutant did not (**Fig S4B**). Thus, direct binding between TAX1BP1 and the TIR motif of NBR1 is required for synergistic functions of NBR1 and TAX1BP1 across multiple in vivo models.

Finally, we explored the specific role of TAX1BP1 in mediating NBR1 flux. Previous studies have reported that TAX1BP1 recruits FIP200 to p62 condensates via NBR1 (Ohnstad *et al*., 2020; Turco *et al*., 2021). To test whether TAX1BP1 makes additional contributions *in vivo*, we co-transfected PentaKO HeLa cells with tf-NBR1 and BFP-TAX1BP1 variants, while monitoring NBR1 flux. Previous reports indicate NBR1 flux is dependent on its LIR motif (Y732A) (Kirkin, Lamark, Sou, *et al*., 2009; Waters *et al*., 2009). Intriguingly, we find that this defect is rescued by wild-type TAX1BP1 (**Fig 6F**). Based on this observation, we hypothesized that TAX1BP1 provides additional LIR motifs to facilitate flux. We then mutated the TAX1BP1 LIR motif, V143S (Tumbarello *et al*., 2015), and assessed synergy with NBR1. Notably, BFP-TAX1BP1^V143S^ neither enhances tf-NBR1 flux, nor rescues tf-NBR1^Y732A^ flux (**Fig 6F**). These data show that beyond recruiting FIP200, TAX1BP1-mediated enhancement of NBR1 flux is supported through the addition of high valency interactions with Atg8-family proteins.

## DISCUSSION

Prior studies have demonstrated that NBR1, TAX1BP1, and p62 play distinct roles in aggrephagy (Ohnstad *et al*., 2020; Sarraf *et al*., 2020; Turco *et al*., 2021). Here, we use peptide-based arrays and cell-based reconstitution to provide additional insights into their cooperative dynamics within cellular autophagy processes. While SQSTM1-like receptors (SLRs) are each independently capable of targeting to lysosomes, we find that their simultaneous expression typically results in competition (**Fig 6C**). A similar phenomenon was recently reported for Zellweger spectrum disorder and neurodegenerative disease, where disease-induced cargoes outcompete constitutive cargo, leading to systemic autophagy failure due to cargo overload (Germain *et al*., 2024). However, in the case of NBR1 and TAX1BP1, we find that heterotypic interactions enable cooperation rather than competition, potentially reducing the burden on cells facing heavy autophagy demands.

Interestingly, our study did not observe the previously reported synergy between p62 and NBR1 (Sánchez-Martín *et al*., 2020; Turco *et al*., 2021). This is perhaps not unexpected. First, biochemical studies have highlighted the importance of proper stoichiometry between p62 and NBR1 (Turco *et al*., 2021). In our cell-based reconstitutions, where receptor titration through co-transfection cannot be precisely controlled, the required stoichiometry is unlikely to be consistently achieved. Alternatively, a key function of p62 is the generation of ubiquitinated condensates, presumed autophagy cargo (Komatsu *et al*., 2010; Sun *et al*., 2018; Zaffagnini *et al*., 2018; Sánchez-Martín *et al*., 2020; Turco *et al*., 2021; Kurusu, Morishita and Komatsu, 2023). Our data suggest that p62’s role in coalescing ubiquitinated cargo, while critical for cargo incorporation and clearance, is not required for NBR1 flux. However, NBR1 could then accelerate the degradation of p62-bound cargo by linking it to its own rapid turnover, a process further expedited by TAX1BP1. This model positions NBR1, one of the earliest conserved autophagy receptors (Kraft, Peter and Hofmann, 2010; Rasmussen *et al*., 2022), as a central orchestrator in this network, with evolutionary adaptors (i.e. p62 and TAX1BP1) subsequently enhancing its functionality and minimizing competition from other autophagy processes.

More broadly, this study presents the first systematic analysis of over 100 autophagy-related SLiMs, evaluating the binding correlation between LC3A, GABARAPL1, FIP200, and TAX1BP1 (**Fig 2A**). While multiple studies have previously detailed the binding determinants within individual SLiMs, our comprehensive approach allows us to extract insights into the binding characteristics of these motifs across a wide array of autophagy-related proteins, thereby allowing us to expand and generalize insights from previous studies.

The utility of our approach is highlighted by comparing our data on NBR1 with the 100 LIR array (**Fig. 2A**). Our structural modeling of the NBR1 SLiM finds residues that facilitate LC3A binding also make substantial contacts in FIP200 binding, possibly suggesting these factors recognize similar overlapping sequences (**Fig S3A-G**). However, our mutagenic peptide array reveals significant differences between FIP200 binding and LC3A binding. Most notably, individual point mutants within the core LIR (732-735) were insufficient to disrupt FIP200 binding (**Fig 3B**). This suggests that FIP200^Claw^ may be more tolerant than ATG8 family members regarding the spacing and size of hydrophobic residues within the motif. Alternatively, the NBR1 LIR contains an extended hydrophobic stretch (YIIIL) that may accommodate different binding modes in the presence of single point mutations. Correspondingly, we were unable to identify FIP200-specific mutations to determine the contribution of the NBR1:FIP200^Claw^ interaction *in vivo*. However, analyzing across >100 LIRs and FIRs, we similarly find discordance between the binding of FIP200 and Atg8-family proteins (**Fig 2C**). This result further highlights that LC3A binding and FIP200 binding are not inseparable events. Rather, multifunctional binding sites likely represent an evolutionary tradeoff between multiple related, but nonidentical, consensus motifs.

What advantage is gained by such close co-evolution of LIRs and FIRs? As previously proposed (Turco *et al*., 2019; Zhou *et al*., 2021), overlapping FIRs and LIRs with differential affinities suggests an intriguing mechanism to drive the sequential recruitment of autophagosomal factors—FIP200 first, followed by the higher affinity Atg8s. This arrangement could ensure that once cargo is bound FIP200 is displaced to prevent its degradation, thus facilitating continuous autophagosome formation. Building on this model, we note that FIP200 binding sites correlate notably better with LC3A binding than with GABARAPL1 binding (**Fig 2C**). From this, we propose that a previously unanticipated functional distinction between the LC3 and GABARAP subfamilies of Atg8 could be a selective role for LC3-family proteins in displacing FIP200 during autophagosome maturation.

The effects of phosphorylation at strategic sites within these motifs add yet another layer of complexity. Phosphorylation positively correlates with both FIP200 and Atg8-family binding, preventing us from concluding a universal sequence of events that is governed in a phospho-dependent manner (**Fig 4B**). Moreover, while phosphomimetics at positions −2 or −1 generally enhance binding, this effect is not universal. Therefore, although a general trend, the enhancement effect of phosphorylation might be selective to certain motifs within the LIR and FIR landscape, and not uniformly applicable across all interactions. At the same time, geometry of the phosphomimetic residues is known to be a suboptimal proxy for phosphorylation (Kliche *et al*., 2022; Popelka and Klionsky, 2022). As such, the role of phosphorylation is likely to be underestimated by our approach. In addition, the role of phosphorylation at other sites was not formally tested. However, the contribution of other acidic residues to the interaction of NBR1 with LC3 and FIP200 is consistent with a broad ability of phosphorylation to affect LIR binding (Rogov *et al*., 2023). This likely enables a more nuanced control over these interactions, which may be critical for the dynamic regulation of autophagy under varying physiological conditions.

Finally, the overlap in binding sites for TAX1BP1, LC3A, and FIP200 introduce an additional complexity (**Fig 1C** and **1D**). Our AlphaFold modeling of NBR1 and TAX1BP1 suggests that the frequent interaction of TAX1BP1 across the LIR peptide array might be explained by the aromatic residues common in all LIRs, which fit into the hydrophobic pocket of TAX1BP1^CC2^ (**Fig 5B**). Consequently, we anticipate the number of in vivo relevant TIR-containing peptides to be overestimated by our peptide array. More comprehensive studies will be necessary to prioritize these sites and clarify their functional significance in vivo. However, an overarching concept that emerges from this study is the potential for LC3-interacting regions to serve as protein interaction hot-spots. This idea further expands the known role of LIRs in autophagy and introduces many new avenues for exploration. Compellingly, a noncanonical AIM – termed the shuffled AIM (sAIM) – has been shown to facilitate interaction with both Atg8-family proteins and another ubiquitin-like protein, UFM1 (Picchianti *et al*., 2023). We anticipate further instances where additional interacting proteins bind to LIRs with significant effects. Investigating these possibilities will be essential for gaining deeper insights into autophagy regulation and its implications for cellular function and disease pathology.

An enduring question is why multifunctional autophagy SLiMs have evolved rather than separate, non-overlapping binding sites. Computational simulations suggest that heterotypic (one-to-many) interactions enhance the efficiency of phase transitions within cellular condensates by lowering the phase boundary considerably compared to a one-to-one binding model (Riback *et al*., 2020; Krishnan *et al*., 2022). This suggests that heterogeneous systems, like those involving the heterotypic SLiM of NBR1, can undergo phase transitions at reduced valency, highlighting a possible evolutionary benefit of receptor cooperation. Future studies will need to fully test these hypotheses. It also suggests that, while each SLR contains sufficient utility to flux on their own, by combining their varied multi-valencies, cells can optimize clearance of targeted cargo (Sun *et al*., 2018; Jakobi *et al*., 2020; Yamasaki *et al*., 2020; Agudo-Canalejo *et al*., 2021). We note that TAX1BP1’s cooperative impact hinges on its LC3 binding capacity (**Fig 6F**). This points to LC3 valency as a critical factor, beyond the previously emphasized role of TAX1BP1 in FIP200 recruitment (Ohnstad *et al*., 2020; Turco *et al*., 2021; Zhang *et al*., 2024). Therefore, the multifunctional SLiM of NBR1, coupled with TAX1BP1’s diverse binding contributions, underscores the role of multi-valency in optimizing cellular clearance mechanisms, reinforcing our understanding of autophagy receptor function and interaction as a complex, cooperative system that enhances adaptability and efficiency.

In conclusion, our study offers new insights into the interactions between autophagy-related SLiMs and their numerous overlapping binding partners. Our findings highlight the diverse and partially overlapping binding motifs of NBR1, illustrating an intricate co-evolutionary balance that fulfills the requirements of three distinct binding events, each necessitating a unique conformation of the unstructured NBR1^725-749^ SLiM. This one-to-many binding modality unveils a complex, cooperative mechanism that enhances cellular adaptability and efficiency. Such cooperation between autophagy receptors and other autophagy factors furthers the mechanistic basis for the widespread role of multifunctional SLiMs in autophagy. These insights lay the groundwork for future research into the molecular mechanisms driving autophagy, potentially leading to novel therapeutic approaches for diseases associated with autophagy dysregulation.

## Supporting information

Supplemental Table 1 - Peptides

## MATERIALS AND METHODS

### Antibodies

All immunoblotting (IB) primary antibodies were diluted 1:1,000 unless otherwise noted. All secondary antibodies were diluted 1:10,000 unless otherwise noted. The following antibodies were used: mouse anti-myc (M4439, Sigma), rabbit anti-V5 (13202, CST), mouse anti-V5 (MCA1360, Biorad), rabbit anti-GST (2625T, CST), rabbit anti-MBP-tag (15089-1-AP, Proteintech), mouse anti-HA (901501, BioLegend), rabbit anti-TAX1BP1 (5105, CST), rabbit anti-LC3A/B (12741S, CST), goat anti-GST-HRP conjugate (RPN1236, Cytiva), goat anti-Rb-HRP conjugate (1706515, Bio-rad), goat anti-Ms-HRP conjugate (1706516, Bio-rad).

### Chemicals and reagents

The following chemicals and reagents were used in this study: 2-mercaptoethanol (BME) (M6250-100ML, Sigma), 41-µm nylon mesh (B0015GZDEQ, Amazon), agar (A10752, Alpha Aesar), agarose (16500500, Thermo Fisher), Amicon Ultra-15 Centrifugal Filters (UFC903024, Sigma), ampicillin (A9518-25G,Sigma), Bafilomycin A1 (11038, Caymen chemical), BCA assay (23227, Pierce), blasticidin (ant-bl1, Invivogen), cOmplete protease inhibitor tablet (5056489001, SIGMA), Dynabeads protein G (10004D, ThermoFisher), EDTA (EDS-500G, Sigma), Glutathione Sepharose 4B resin (95016-984, VWR), glycerol (G2025-1L, Sigma), HEPES (H3375-1KG, Sigma), Imidazole (O3196-500, FischerSci), Intercept^TM^ (TBS) Blocking Buffer (927-60003, LI-COR), Isopropyl-β-D-thiogalactopyranoside (IPTG) (BP1755-10, Fischersci), Lipofectamine 3000 (97064-850, FischerSci), Novex 4-20% Tris-Glycine gels (WXP42020BOX, Thermo Fisher), Phusion High-Fidelity DNA polymerase (M0530L, NEB), Pico PLUS chemiluminescent substrate (PI34577, FischerSci), polybrene (H9268-5G, Sigma), potassium chloride (P217-500, FisherSci), puromycin (ant-pr-1, Invivogen), PVDF membranes 0.2 µm (#ISEQ00010, Sigma), Sodium Phosphate Dibasic Monohydrate (S373-500, FisherSci), Sodium Phosphate Monobasic Monohydrate (S369-1, FischerSci), sodium chloride (6438, FisherSci), sodium dodecyl sulfate (SDS) (74255-250G, Sigma), sucrose (BP220-1, FisherSci), Taq DNA ligase (M0208L, NEB), Tris(2-carboxyethyl)phosphine hydrochloride (TCEP) (97064-850, VWR), Tris base (T1378-5KG, Sigma), TritonX-100 (T9284500ML, Sigma), tryptone (DF0123-17-3, FisherSci), Tween-20 (BP337500, FisherSci), T5 exonuclease (M0363S, NEB), urea (U15-3, Fischersci), yeast extract (BP1422-2, FisherSci), zeocin (ant-zn-1, Invivogen),

### Vectors

All plasmids are listed in Supplementary Table 2.

### Isothermal assembly

PCR inserts were amplified with Phusion High-Fidelity DNA Polymerase (M0530L, NEB). Amplification primers were designed with a 30 bp overlap with the linear ends of restriction-digested vectors. Linearized vectors were dephosphorylated using calf intestinal phosphatase (M0290S, NEB). All inserts and vectors were purified using 0.9% agarose gel prior to isothermal assembly (D4002, Zymo Research). 50 ng of linearized vector DNA was mixed with an isomolar amount of purified insert(s). 2.5 µl of DNA mix was incubated with 7.5 µl isothermal assembly master mix at 50°C for 20 minutes. Product of the isothermal assembly reaction was transformed into NEB Stable cells (C3040H, NEB). Transformed cells were plated on 1.5% agar plates with LB media (10 g/l tryptone, 5 g/l yeast extract, 5 g/l NaCl). All cultures and plates were grown at 34°C overnight and included 100 µg/ml ampicillin. Overnight cultures were pelleted at 3,000 *g* for 10 min and plasmid DNA was extracted and purified using a Qiagen miniprep kit (27106, Qiagen). Sequences were verified by Sanger sequencing (Eton Bioscience Inc).

### Tissue Culture

K562 ATG7^KO^/TAX1BP1^KO^ cells expressing tf-NBR1 were generated previously (Ohnstad *et al*., 2020). HeLa PentaKO cells were a gift from Dr. Richard Youle (Lazarou *et al*., 2015). All cells were grown in a standard water jacketed incubator with 5% CO_2_. K562 cells were grown in IMDM media (10-016-CV, Corning) with 10% FBS (26140079, Thermo Fisher) and 1x pen/strep (15140122, Thermo Fisher). HEK293T and HeLa PentaKO cells were grown in DMEM media (10-013-CV, Thermo Fisher) with 10% FBS and 1x pen/strep. All cells were passaged < 25 times. For passaging, cells were washed with HBSS (14025092, Thermo fisher) and trypsinized with Trypsin-EDTA (25300-054, Gibco). Puromycin (2 µg/ml), blasticidin (5 µg/ml), and zeocin (50 µg/ml) were added when necessary for selection.

### Transient transfection and nucleofection

HEK293T cells with grown overnight in Opti-MEM Reduced Serum media with 5% FBS (51985-034, Thermo Fisher) to 90% confluence. HeLa PentaKO cells were grown overnight in Opti-MEM Reduced Serum media with 5% FBS to 70% confluence. Cells were transfected using Lipofectamine 3000 reagent (L3000008, Life Technologies) according to manufacturer’s recommendations. K562 cells were nucleofected using a Nucleofector^TM^ 2b Device (Lonza) using nucleofector kit T (VACA-1002, Lonza) and protocol T-016.

### Immunoprecipitation

Cells were collected and resuspended in lysis/IP buffer (50 mM HEPES pH 7.4, 150 mM NaCl, 2 mM EDTA, 1% Triton X-100, 2x cOmplete protease inhibitor tablet [5056489001, SIGMA]) or IP Buffer with 0.5 mM DTT. Cells were incubated on ice for 20 min and pelleted at 1000 *g* 10 min at 4°C. Supernatant was normalized by total protein using BCA assay (23227, Pierce) prior to IP. Normalized extract was applied to Glutathione Sepharose 4B resin (95016-984, VWR) pre-bound to purified proteins, or protein G dynabeads (10004D, Thermo Fisher) pre-bound to the indicated antibodies. Incubations were allowed to proceed for 1 h at 4°C. Beads were washed 3x times with one tube change. Proteins were eluted by boiling at 70°C in 1x Laemmli Loading Buffer (3x stock: 189 mM Tris pH 6.8, 30% glycerol, 6% SDS, 10% beta-mercaptoethanol, bromophenol blue).

### Gel electrophoresis and immunoblotting

Gel electrophoresis was performed at 195V for 65 min in Novex 4-20% Tris-Glycine gels (WXP42020BOX, Thermo Fisher). For western blotting, samples were transferred for 60 min to 0.2 µm PVDF membranes (#ISEQ00010, Sigma) using a Semi-dry transfer cell (Bio-Rad). Membranes were blocked for 20 min in Intercept^TM^ (TBS) Blocking Buffer (927-60003, LI-COR). Primary antibodies were incubated overnight at 4°C. Blots were rinsed 3x 5 min in TBS-T (10mM Tris pH 7.9, 150 mM NaCl, 0.05% Tween-20, 0.25 mM EDTA). Secondary antibodies (1:10,000 dilution in Intercept^TM^ [TBS] Blocking Buffer with 0.2% Tween-20 and 0.01% SDS) were incubated for 1 h at room temperature. Blots were rinsed 3x 10 min in TBS-T and imaged using fluorescence (LI-COR Odyssey CLx Imager).

### Flow cytometry

Adherent cells were trypsinized, pelleted, washed in cold HBSS, and filtered through 41-µm nylon mesh (B0015GZDEQ, Amazon) prior to analysis. K562 cells were washed in cold HBSS and filtered through 41-µm nylon mesh prior to analysis. Data were collected on a Cytoflex S Cytometer (Beckman). Data were analyzed using FlowJo v 10 (FlowJo, LLC) and R. The red:green ratios of tf-labelled reporters were normalized to a Bafilomycin treated control where applicable.

### Statistical analysis

All statistical analysis was performed using Prism 10 (GraphPad). All statistical tests are indicated in the relevant figure legends. All tests were two-tailed with *p*< 0.05 as the minimum threshold for statistical significance. Prior to the indicated tests, Shapiro-Wilk was used to test normality. The Brown-Forsythe test was used to test for unequal variance for ANOVA. ANOVA with multiple comparison used Dunnet T3 correction or Tukey correction as appropriate. Number of replicates (*n*) used for each test are indicated in the relevant figure legend.

### AlphaFold structural modeling

Sequences were obtained from Uniprot and the LIR-central database (Chatzichristofi *et al*., 2023). AF2-multimer was used with MMseqs2 through Google Colab (<u>htps://colab.</u> <u>research.google.com/github/sokrypton/ColabFold/blob/main/AlphaFold2.ipynb#scrollTo=UGUB LzB3C6WN</u>). The top ranked structure was relaxed using AMBER and examined using Pymol V2.4.1 (PyMOL V2.4.1, Schrödinger, LLC). Predicted LDDT (pLDDT) per residue was estimated by pLDDT metric output by AF2 as a metric of model confidence. pLDDT scores were colored in pymol using the Pymol extension Color AlphaFold (https://github.com/cbalbin-bio/pymol-color-alphafold).

### Buried surface area calculations

The buried surface area (BSA) of NBR1 amino acids upon binding to different binding partners was calculated using the Proteins, Interfaces, Structures and Assemblies (PISA) server at the European Bioinformatics Institute (http://www.ebi.ac.uk/pdbe/prot_int/pistart.html). Each relaxed model from the AlphaFold structural modeling was analyzed for the BSA area at the interface of NBR1 and its binding partner.

### Peptide array design

Peptides arrays were printed by the MIT Biopolymers & Proteomics Core. LIR sequences were drawn from the LIR-Central database (Chatzichristofi *et al*., 2023) (**Supplemental Table 1**). FIR sequences were drawn from (Fujita *et al*., 2013; Smith *et al*., 2018; Ravenhill *et al*., 2019; Fu *et al*., 2021; Zhou *et al*., 2021). The NBR1 sliding frame array was printed using 20 aa sections of NBR1^711-759^ printed with a 1 aa frame shift per spot. Sequences containing serine or threonine residues at −2 or −1 positions relative to the LIR were mutated to aspartic acid and printed in parallel. The NBR1 mutational peptide array was printed using NBR1^725-749^, varying each position by all possible residues. As binding controls, the top two rows and botom row were printed with wild type peptide.

### Peptide array probing

Peptide arrays were reconstituted in methanol and sonicated briefly (15-30 s) prior to washing 3x 5 min in TBS-T (10mM Tris pH 7.9, 150 mM NaCl, 0.05% Tween-20, 0.25 mM EDTA), followed by a 1 h incubation in TBS-T + 0.5 mM TCEP to fully reduce peptide spots. Arrays were blocked in Intercept^TM^ + 0.2% Tween-20 + 0.5 mM TCEP overnight at 4°C. Arrays were then stripped prior to first probe and between subsequent probes. Arrays were stripped by washing 3x 10 min in H_2_O, sonicating at 40°C 3x 10 min in stripping mix A (8M Urea, 2% SDS, 0.7% BME, in PBS pH 7), washing 3x 10 min in stripping mix B (10% acetic acid, 50% ethanol, 40% H_2_O), re-wetting in alcohol 3x 10 min with brief sonication (15-30 s), washing 3x 10 min in TBS-T + 0.5 mM TCEP, and blocking in Intercept^TM^ + 0.2% Tween-20 + 0.5 mM TCEP 1 H at 4°C.

Peptide arrays were probed with 100 nM of each indicated protein in Intercept^TM^ + 0.2% Tween-20 + 0.5 mM TCEP for 2 h at RT (FIP200^Claw^ was probed in Intercept^TM^ + 0.5% Triton X-100 + 0.5 mM TCEP for 2 h at RT). Probed membranes were washed 3x 10 min in TBS-T prior to probing with primary antibodies as indicated in TBS-T (10mM Tris pH 7.9, 150 mM NaCl, 0.2% Tween-20, 0.25 mM EDTA) with 5% milk powder overnight at 4°C. Probed arrays were washed 3x 10 min in TBS-T followed by incubation with secondary HRP conjugated antibodies as indicated in TBS-T + 5% milk for 1 h at 4°C then washed 3x 10 min in TBS-T.

HRP probed peptide arrays were visualized with Pico PLUS chemiluminescent substrate (PI34577, FischerSci) according to the manufacture’s recommendations using a Syngene imager. Images were analyzed and quantified using ImageLab 6.0 (BioRad). Images were scored blindly to determine a bind/no bind cutoff. In the LIR family array, peptide spots that failed to bind to either LC3A, GABARAPL1, FIP200^Claw^ or TAX1BP1^CC2^ were designated as “never-binders” and were excluded from further analysis.

### Fluorescence anisotropy

FITC-NBR1 peptide was sourced from Biomatik and reconstituted in DMSO before dilution to Phosphate buffered SEC buffer (0.05M NaH2PO4 pH 7.4 pH, 100 mM NaCl, 0.5 mM TCEP). Purified proteins GST-LC3A, GST-FIP200^Claw^, and GST-TAX1BP1^CC2^-MBP were purified as described and serially diluted in Phosphate buffered SEC buffer (0.05M NaH2PO4 pH 7.4 pH, 100 mM NaCl, 0.5 mM TCEP) with 50 nM FITC-NBR1^725-749^. Assays were performed on a Synergy Neo2 plate reader (BioTek). Binding curves were fit using a One Site: Total nonlinear regression model to determine K_D_.

### Protein expression and purification

GST-tagged proteins were expressed in E.coli Roseta 2 cells overnight at 16 °C with 0.4 mM IPTG and purified using a HisTrap HP 5 ml column (Cytiva) equilibrated in (75 mM Tris pH 8, 500 mM NaCl, 5 mM BME, 20 mM imidazole, 10% glycerol) and eluted in (75 mM Trsi pH 8, 100 mM NaCl, 5 mM BME, 400 mM imidazole, 10% glycerol) via a stepwise imidazole gradient. The protein was applied to a HiLoad 26/600 Superdex 200 column (Cytiva) equilibrated in (0.05 M NaH_2_PO_4_ pH 7.4, 100 mM NaCl, 5 mM BME). Fractions containing purified protein were concentrated, frozen using liquid nitrogen, and stored at −80°C.

HA-tagged MAP1LC3A and GABARAPL1 were expressed in E.coli Roseta 2 cells overnight at 16 °C with 0.4 mM IPTG and purified using HisTrap HP 5 ml column (Cytiva) equilibrated in (75 mM Tris pH 8, 500 mM NaCl, 5 mM BME, 20 mM imidazole, 10% glycerol) and eluted in (75 mM Trsi pH 8, 100 mM NaCl, 5 mM BME, 400 mM imidazole, 10% glycerol) via a stepwise imidazole gradient. Eluted protein was dialyzed to (20 mM Hepes pH 7.4, 150 mM NaCl, 5 mM BME, 5% glycerol) and cleaved with TEV protease overnight at 4 °C. Cleaved and dialyzed protein was applied to a HiLoad 26/600 Superdex 200 equilibrated in (20 mM Hepes pH 7.4, 150 mM NaCl, 5 mM BME, 5% glycerol). Fractions containing purified protein were concentrated, frozen using liquid nitrogen, and stored at −80°C.

---

**Figure S1.**
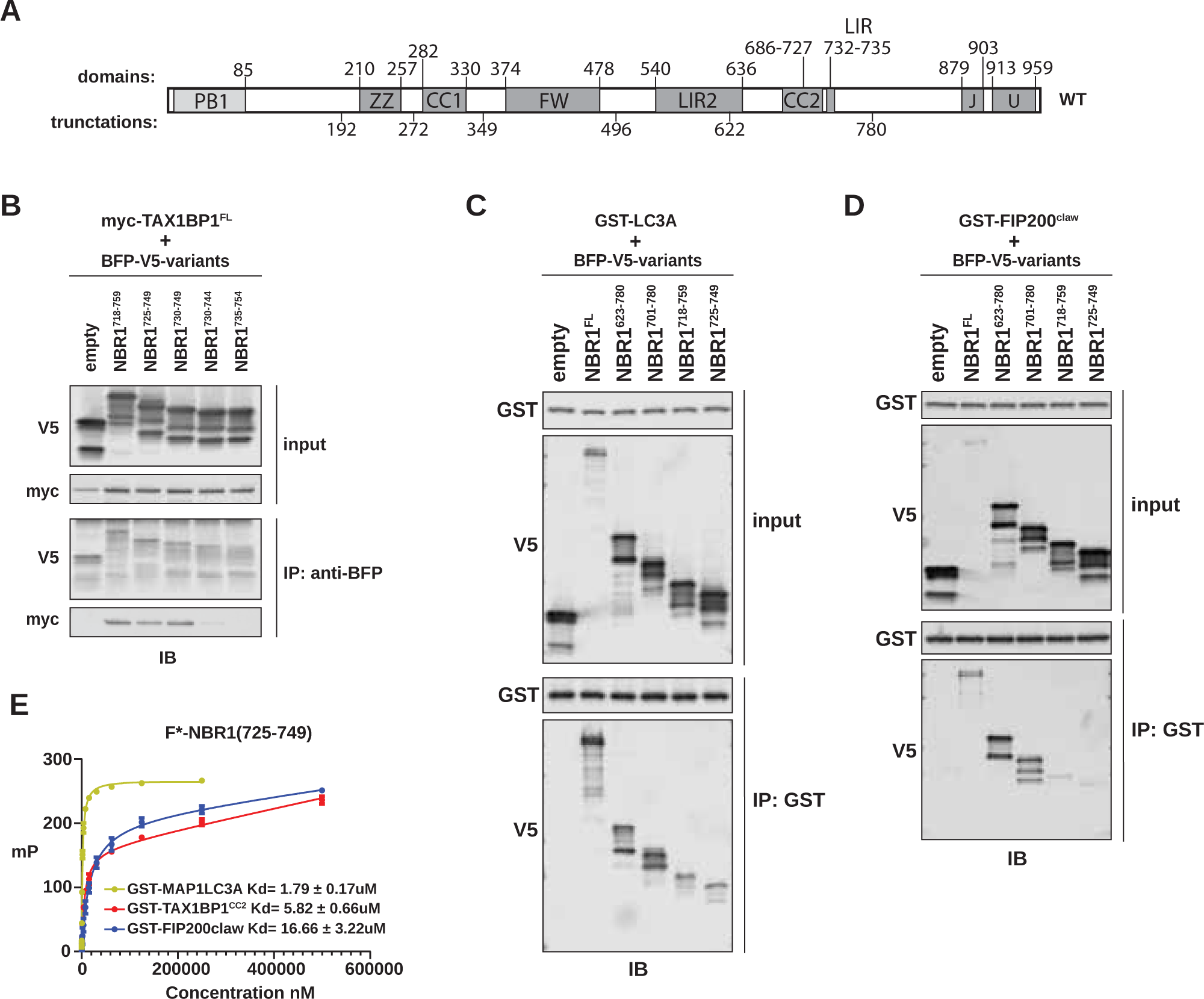
NBR1 binds LC3A, FIP200^Claw^, and TAX1BP1^CC2^ through a shared unstructured region. **A)** Schematic of NBR1 domain architecture. Domain boundaries are listed above the schematic. Truncation boundaries are listed below the schematic. **B)** HEK293T cells were co-transfected with full-length (FL) myc-TAX1BP1 and the indicated BFP-V5-NBR1 truncations. Extracts derived from transfected cells were immunoprecipitated (IP) with protein G dynabeads conjugated with anti-BFP antibody. Input and eluates were resolved by SDS-PAGE followed by immunoblotting (IB) with the indicated antibodies. **C, D)** HEK293T cells were transfected with indicated BFP-V5-NBR1 truncations. Extracts derived from transfected cells were incubated with GST-resin pre-bound with purified GST-MAP1LC3A or GST-FIP200^Claw^. Input and eluates were resolved by SDS-PAGE followed by immunoblotting (IB) with the indicated antibodies. **E)** Purified GST-MAP1LC3A, GST-FIP200^Claw^, or GST-TAX1BP1^CC2^ were serially diluted and incubated with 50 nM FITC-labelled NBR1^725-749^. Fluorescence anisotropy was measured by plate reader and analyzed to determine K_D_.

**Figure S2.**
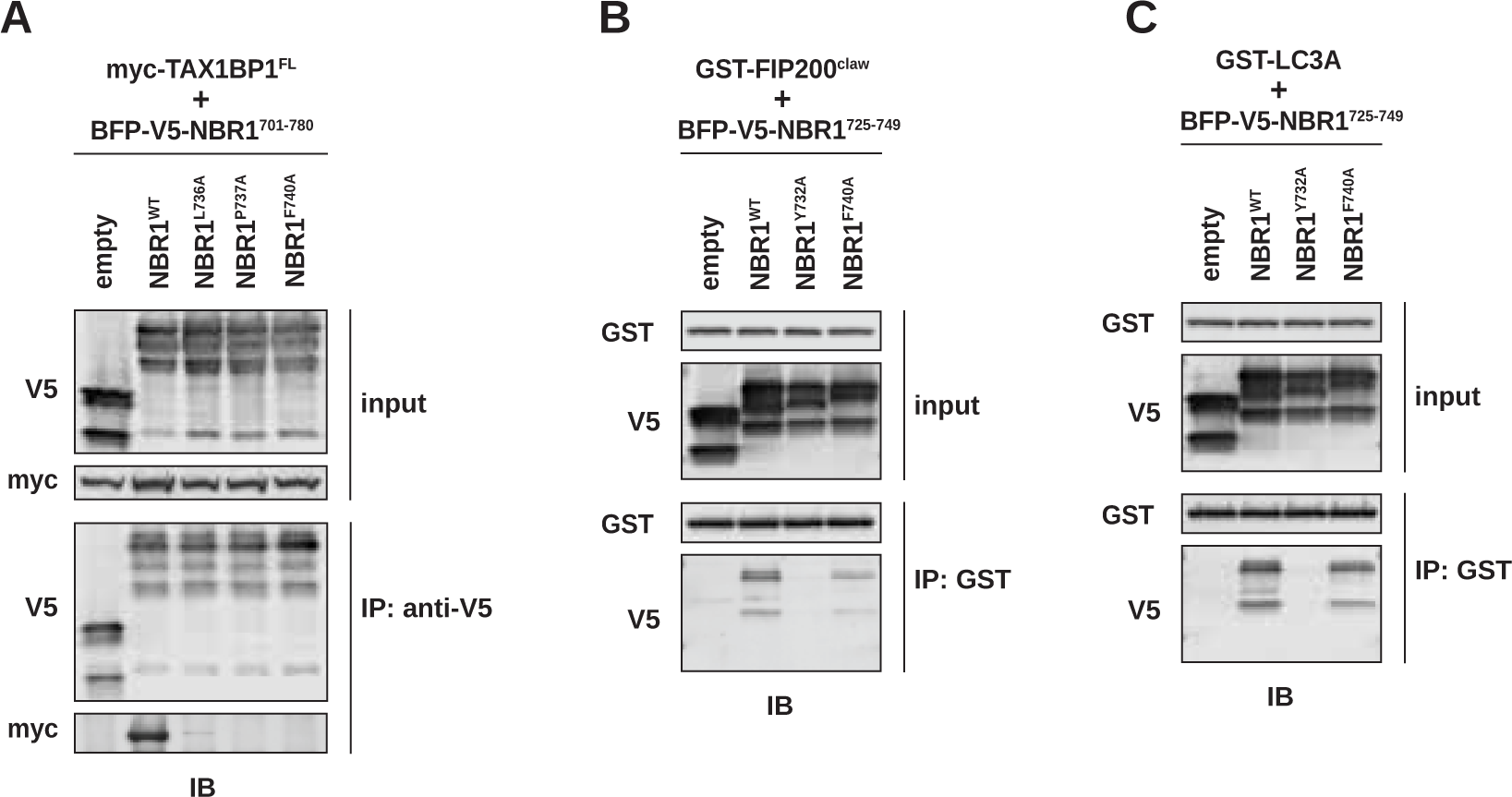
Selective mutations disrupt NBR1 interaction with LC3 or TAX1BP1^CC2^. **A)** HEK293T cells were co-transfected with full-length (FL) myc-TAX1BP1 and indicated BFP-V5-NBR1 variants. Extracts derived from transfected cells were immunoprecipitated (IP) with protein G dynabeads conjugated with anti-V5 antibody. Input and eluates were resolved by SDS-PAGE followed by immunoblotting (IB) with the indicated antibodies **B, C)** HEK293T cells were transfected with indicated BFP-V5-NBR1 variants. Extracts derived from transfected cells were incubated with GST-resin pre-bound with purified GST-FIP200^Claw^(B) or GST-MAP1LC3A (C). Input and eluates were resolved by SDS-PAGE followed by immunoblotting (IB) with the indicated antibodies.

**Figure S3.**
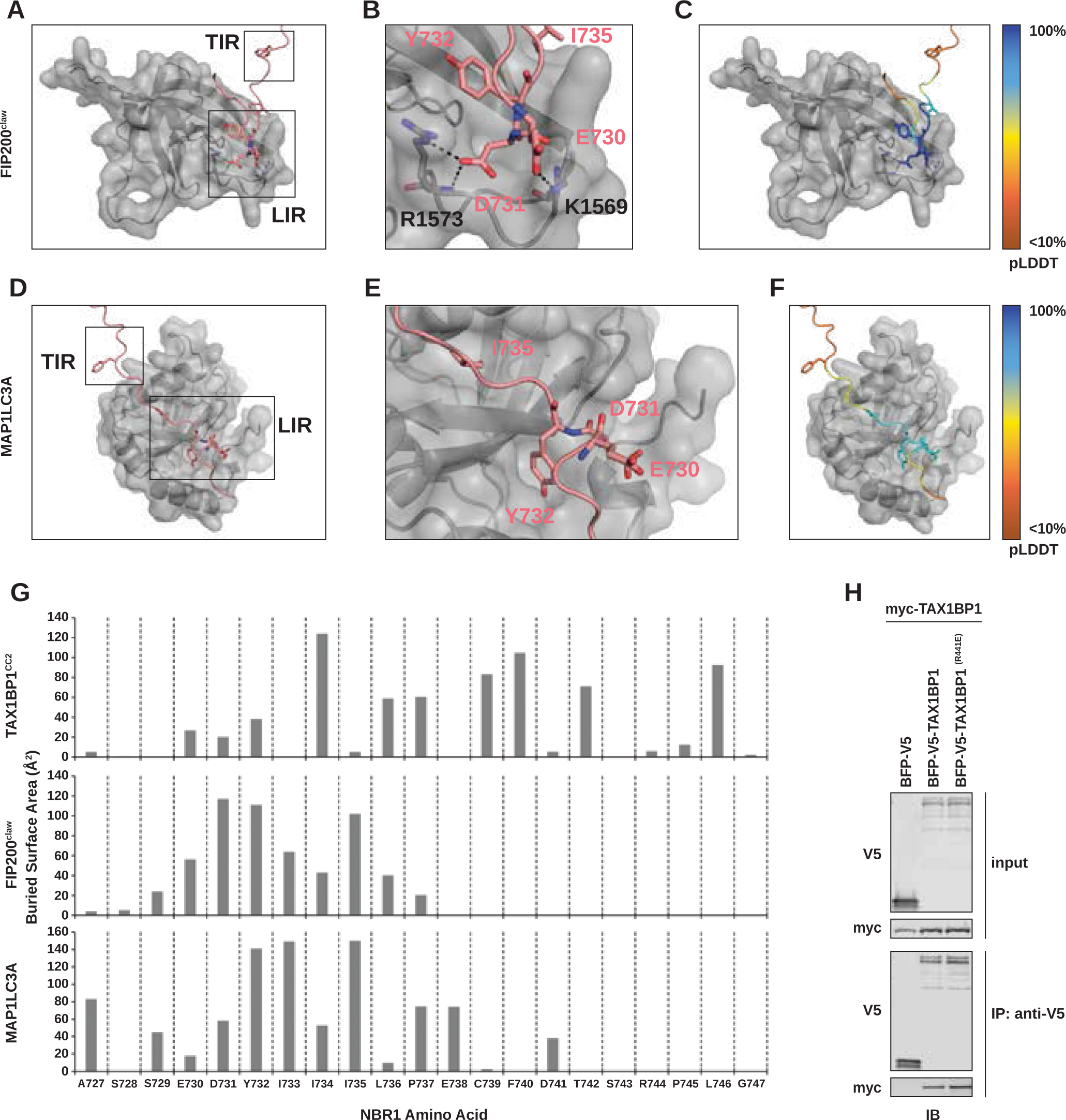
Comparative interactions between NBR1 and LC3, FIP200^Claw^, and TAX1BP1^CC2^. **A)** Predictive structural model of NBR1^725-749^ with FIP200^Claw^. Model was generated using AlphaFold AF2-multimer with MMseqs2 through Google Colab. TIR inset shows no meaningful interactions. **B)** LIR inset from **A**, highlighting core LIR residues (NBR1^Y732^ and NBR1^I735^) and the interaction between NBR1^D731^ and NBR1^E730^ with FIP200^R1573^ and FIP200^K1569^, respectively. **C)** Predicted IDDT per position (pLDDT) for NBR1^725-749^ in complex with FIP200^Claw^. Blue is high confidence, orange is low confidence. **D)** Predictive structural model of NBR1^725-749^ with MAP1LC3A. Model was generated using AlphaFold AF2-multimer with MMseqs2 through Google Colab. TIR inset shows no meaningful interactions. **E)** LIR inset from **D**, highlighting core LIR residues (NBR1^Y732^ and NBR1^I735^) and NBR1^D731^ and NBR1^E730^. **F)** Predicted IDDT per position (pLDDT) for NBR1^725-749^ in complex with MAP1LC3A. Blue is high confidence, orange is low confidence. **G)** Buried surface area (BSA) analysis using the PDBePISA server of NBR1^725-749^ with MAP1LC3A, FIP200^Claw^, and TAX1BP1^CC2^. NBR1 residues spanning 727-747 are shown. **H)** HEK293T cells were co-transfected with full-length (FL) myc-TAX1BP1 and indicated BFP-V5-TAX1BP1 variants. Extracts derived from transfected cells were immunoprecipitated (IP) with protein G dynabeads conjugated with anti-V5 antibody. Input and eluates were resolved by SDS-PAGE followed by immunoblotting (IB) with the indicated antibodies.

**Figure S4.**
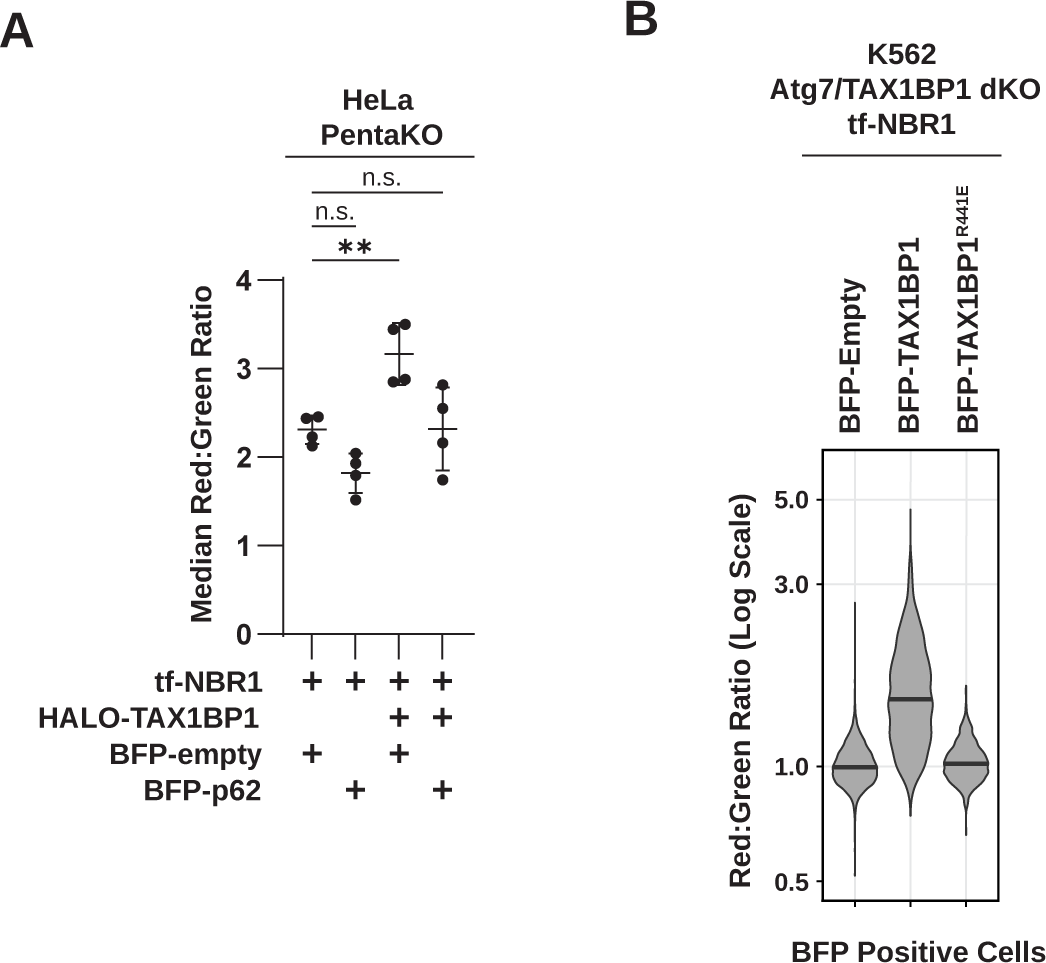
Synergy between NBR1 and TAX1BP1. **A)** HeLa PentaKO cells were co-transfected with tf-NBR1, HALO-TAX1BP1, and the indicated BFP-tagged receptors. Triple-positive cells were analyzed by flow cytometry with red:green ratio as a readout of NBR1 flux. Graphs represent mean +/- SD from three independent experiments. *n* > 10,000 cells per sample. *p* values were determined using a one-way ANOVA (*p* = 0.0006). Multiple comparisons were made using Dunnet T3 correction. **, *p* < 0.01, *ns*, not significant. **B)** *Atg7^KO^/TAX1BP1^KO^* K562 cells expressing tf-NBR1 were electroporated with the indicated BFP-tagged variants of TAX1BP1. BFP expressing cells were analyzed by red:green ratio as a readout for flux. *n* > 10,000 cells per sample.

**Supplemental Table 2.** Vectors used in this study.

|  |  |
| --- | --- |
| pET16bMT-01 | (69662-3, EMD Millipore) |
| pET16bMT (His-TS-TEV-FLAG-HA)-MAP1LC3A (truncated form) | This study |
| pET16bMT (His-TS-TEV-FLAG-HA)-GABARAPL1 (truncated form) | This study |
| pGEX-KT (GST-thrombin-His6-TEV) | Dartmouth Biomolecular Targeting core (77331, ATCC) |
| pGEX-KT (GST-thrombin-His6-TEV)-MAP1LC3A | This study |
| pGEX-KT (GST-thrombin-His6-TEV)-FIP200 CLAW short (1490-1594) | This study |
| pGEX-KT (GST-thrombin-His6-TEV)-TAX1BP1 (346-506)-3C-MBP | This study |
| pHAGE-myc-AP2-TAX1BP1 | (Shoemaker *et al.*, 2019) |
| pHAGE-myc-AP2-NBR1 | (Shoemaker *et al.*, 2019) |
| pLenti TagBFP-V5 empty | This study |
| pLenti TagBFP-V5 NBR1 | This study |
| pLenti TagBFP-SQSTM1 | This study |
| pLenti TagBFP-V5-NDP52 | This study |
| pLenti TagBFP-HA-OPTN | This study |
| pLenti TagBFP-V5-TAX1BP1 | (Ohnstad *et al.*, 2020) |
| pLenti TagBFP-V5-TAX1BP1 V143S | This study |
| pLenti TagBFP-V5-TAX1BP1(R441E) | This study |
| pLenti TagBFP-V5-NBR1 193-end F | (Ohnstad *et al.*, 2020) |
| pLenti TagBFP-V5-NBR1 273 - end F | (Ohnstad *et al.*, 2020) |
| pLenti TagBFP-V5-NBR1 350 - end F | (Ohnstad *et al.*, 2020) |
| pLenti TagBFP-V5-NBR1 497 - end F | (Ohnstad *et al.*, 2020) |
| pLenti TagBFP-V5-NBR1 623 - end F | (Ohnstad *et al.*, 2020) |
| pLenti TagBFP-V5-NBR1 781 - end F | (Ohnstad *et al.*, 2020) |
| pLenti TagBFP-V5-NBR1 1- 192 R | (Ohnstad *et al.*, 2020) |
| pLenti TagBFP-V5-NBR1 1- 272 R | (Ohnstad *et al.*, 2020) |
| pLenti TagBFP-V5-NBR1 1- 349 R | (Ohnstad *et al.*, 2020) |
| pLenti TagBFP-V5-NBR1 1- 496 R | (Ohnstad *et al.*, 2020) |
| pLenti TagBFP-V5-NBR1 1- 622 R | (Ohnstad *et al.*, 2020) |
| pLenti TagBFP-V5-NBR1 1- 780 R | (Ohnstad *et al.*, 2020) |
| pLenti TagBFP-V5-NBR1 623-780 | This study |
| pLenti TagBFP-V5-NBR1 701-780 | This study |
| pLenti TagBFP-V5-NBR1 718-759 | This study |
| pLenti TagBFP-V5-NBR1 725-749 | This study |
| pLenti TagBFP-V5-NBR1 730-749 | This study |
| pLenti TagBFP-V5-NBR1 730-744 | This study |
| pLenti TagBFP-V5-NBR1 735-754 | This study |
| pLenti TagBFP-V5-NBR1 (Y732A) | This study |
| pLenti TagBFP-V5-NBR1 701-780 (L736A) | This study |
| pLenti TagBFP-V5-NBR1 701-780 (P737A) | This study |
| pLenti TagBFP-V5-NBR1 725-749 (F740A) | This study |
| pLenti TagBFP-V5-NBR1 725-749 (Y732A) | This study |
| pLenti CMV tf-NBR1 | (Shoemaker *et al.*, 2019) |
| pLenti CMV tf-NBR1 (Y732A) | This study |
| pLenti CMV tf-NBR1 (F740A) | This study |
| pLenti CMV tf-V5-TAX1BP1 | (Shoemaker *et al.*, 2019) |
| pLenti CMV tf-V5-TAX1BP1 (R441E) | This study |
| pLenti CMV tf-V5-TAX1BP1 (V143S) | This study |
| pLenti CMV tf-SQSTM1 | (Shoemaker *et al.*, 2019) |
| pLenti CMV tf-NDP52 | (Shoemaker *et al.*, 2019) |
| pLenti CMV tf-OPTN | (Shoemaker *et al.*, 2019) |
| pLenti CMV Halo-V5-TAX1BP1 | This study |

